# Speech Tracking in fMRI Reveals a Distinct Response to Sentence Boundaries During Comprehension

**DOI:** 10.64898/2026.10.13.758938

**Authors:** Orel Levy-David, Refael Tikoschinski, Roi Reichart, Elana Zion Golumbic, Isabelle Deschamps, Galit Agmon

## Abstract

Psycholinguistic research and linguistic theory suggest that comprehending continuous speech involves segmenting it into sentences. Yet, direct neuroimaging evidence for this process remains scarce, partly because sentence boundaries coincide with acoustic, prosodic, lexical, and syntactic changes. Here, we used whole-brain fMRI to test whether sentence boundaries elicit a neural response beyond these accompanying speech properties during spontaneous speech comprehension. We modelled BOLD responses to sentence-final words alongside continuous regressors characterizing accompanying speech properties. This approach revealed sensitivity to continuously varying features of speech across multiple brain regions. Notably, the acoustic envelope predicted BOLD responses in bilateral primary auditory cortices. Responses associated with prosodic breaks and emphasis were found in right temporal cortex, syntactic bottom-up node count in the left superior temporal sulcus, and word concreteness in the parahippocampal gyri. Critically, sentence-final words were associated with an increased activation in right temporal regions, after accounting for acoustic, prosodic, lexical, semantic and syntactic properties. These findings provide evidence for a neural response to sentence boundaries consistent with active sentence segmentation. More generally, this study demonstrates the value of continuous-regressor modelling for investigating speech comprehension with fMRI.

## 1. Introduction

The concept of a sentence is fundamental in language research, yet it remains unclear whether sentences constitute a psychologically relevant unit during continuous speech comprehension. This question is easily overlooked because sentences are largely taken for granted in the way language is usually studied. Syntactic theories typically take the sentence as their basic unit of analysis, laboratory experiments commonly use isolated sentences as stimuli, and most computational models of language are trained on written text, where sentence boundaries are explicitly marked by punctuation. In everyday speech, however, sentences are not presented to the listener as pre-segmented units: rather, speech is continuous, and sentence boundaries must be inferred from the unfolding signal. What evidence do we have that listeners actually segment this continuous signal into sentences?

Early psycholinguistic research suggests that syntactic structure shapes how listeners perceptually organize and remember speech. A series of experiments found that brief clicks tend to be perceptually attracted toward major syntactic boundaries, even when the local acoustic signal is held constant, supporting the view that syntactic structure organizes speech into coherent processing units (Fodor & Bever, 1965, 1965; Garrett et al., 1966). Specific evidence for sentence-level organization comes from Jarvella (1971), who showed that the same word sequence is recalled more accurately when it occurs within a single sentence than when it spans a sentence boundary. Together, these findings support the psychological relevance of syntactic grouping and suggest that sentences may function as coherent cognitive gestalts during speech comprehension.

Sentence boundaries also received considerable attention in reading research. Readers consistently spend more time processing sentence-final words, an effect commonly described as *sentence wrap-up* (Camblin et al., 2007; Just et al., 1982; Just & Carpenter, 1980; Mitchell & Green, 1978; Rayner, 1975; Rayner et al., 1989, 2000). The processes underlying this slowdown remain debated, with proposed explanations including the completion of unresolved interpretive processes and implicit prosody (Hirotani et al., 2006; Stowe et al., 2018; Warren et al., 2009). Sentence-final slowing in reading does not, by itself, distinguish among the processes that may occur at sentence endings, such as boundary detection, syntactic and semantic integration, or implicit prosodic processing. Moreover, since written language typically contains explicit boundary cues (through punctuation or isolated sentence presentation), reading research offers only limited insight into how listeners identify sentence boundaries within continuous speech, where those boundaries must be inferred from the unfolding signal.

Neural responses offer a way to investigate this process, allowing us to study how language unfolds under more naturalistic settings (Hasson & Honey, 2012; Nastase et al., 2020). Continuous naturalistic speech is particularly valuable for investigating sentence segmentation, because listeners must identify boundaries within the ongoing signal, rather than receive sentences as isolated units, as is common in controlled experiments. Isolating a neural response to sentence segmentation, however, also presents a related challenge. As mentioned, boundary-associated responses may reflect syntactic integration, linguistic prediction, or prosodic processing rather than sentence segmentation itself. In speech, acoustic and prosodic properties of the input signal can also elicit neural responses at sentence boundaries. The critical question, therefore, is whether sentence boundaries evoke a neural signature beyond that accounted for by these accompanying properties. Identifying such a signature would support the possibility that the brain treats sentences as important cognitive units, actively marking their completion. If listeners construct sentence-level representations during online comprehension, how does the brain determine where one sentence ends and the next begins?

### 1.1. Sentence-Final Words: Prosodic, Lexical and Syntactic Properties

Listeners can draw on multiple sources of information in order to detect sentence boundaries in speech, some of which are higher-level linguistic expectations while others are more closely tied to the acoustic input. In this section, we review the prosodic, lexical, and syntactic properties associated with sentence boundaries.

Perhaps the most intuitive cue to sentence boundaries is prosody, the patterns of pitch, duration, pausing, intensity, and other acoustic features that unfold over an utterance (de Pijper & Sanderman, 1994; Streeter, 1978). Listeners use these cues to interpret syntactic structure and resolve ambiguities (Beach, 1991; Cutler et al., 1997; Hawthorne & Gerken, 2014; Kjelgaard & Speer, 1999; Klatt, 1975; Lehiste, 1973; Millotte et al., 2007; Morgan, 1996; Price et al., 1991). In particular, prosodic cues for sentence endings include the lengthening of sentence-final words, resetting of pitch and intensity, and pausing before the next sentence (Cooper & Paccia-Cooper, 1980; Ladd, 1988; Strangert, 1992). The contribution of prosody to segmentation is evident even when listeners cannot understand the linguistic content: in unfamiliar or artificial languages, prosodic cues can support the identification of structural units in speech (Langus et al., 2012; Matzinger et al., 2021; Pilon, 1981; Wakefield et al., 1974). Because these acoustic and prosodic properties covary with sentence-final position, identifying a neural response associated with sentence boundaries beyond these properties requires accounting for their contribution to the measured response.

Sentence-final position effects may also be related to word predictability as the sentence unfolds. As linguistic context accumulates, later words become more predictable, possibly peaking at sentence-final positions. Evidence consistent with this pattern comes primarily from reading paradigms: N400 amplitude, which is strongly modulated by word predictability (Kölbl et al., 2026; Kutas & Federmeier, 2011; Michaelov et al., 2024), tends to decrease with increasing word position within a sentence (Dambacher et al., 2006; Kutas & Hillyard, 1983; Payne et al., 2015; Van Petten & Kutas, 1990). More directly, Huizeling et al. (2022) reported a negative correlation between n-gram surprisal and position within the sentence, indicating that later words tend to be less surprising given their preceding context. Lexical predictability is therefore an important potential covariate of sentence-final position.

Lexical prediction, however, is only one component of incremental language comprehension. As speech unfolds, listeners also continuously construct and update syntactic representations (see Coopmans et al., 2026, for a recent review). A growing body of work has shown that measures of incremental syntactic processing explain variation in neural responses beyond that accounted for by lexical surprisal, suggesting that both contribute to the neural processes underlying comprehension (Brennan et al., 2016; Coopmans et al., 2025; Giglio et al., 2024; Lopopolo et al., 2021; Nelson et al., 2017; Shain et al., 2022; Stanojević et al., 2023; Zioga et al., 2023). The evolving syntactic representation may itself provide information about whether a sentence is complete or requires additional material. One measure particularly relevant in this context is *bottom-up node count* (Bhattasali et al., 2019; Brennan et al., 2012, 2016; Cai et al., 2026; Giglio et al., 2024; Gwilliams et al., 2025; Li & Hale, 2019; Lopopolo et al., 2021; Nelson et al., 2017; Reddy & Wehbe, 2021; Stanojević et al., 2023; Weissbart & Martin, 2024), an incremental measure derived from constituency structure that assigns to each word the number of syntactic constituents completed at that word. In English, with its predominantly right-branching structure, multiple constituents often become complete simultaneously at sentence-final words, resulting in relatively high bottom-up node counts at sentence endings (e.g., Coopmans et al., 2026). Neural responses associated with bottom-up node count have been reported primarily in left temporal language regions (Brennan et al., 2016; Giglio et al., 2024; Lopopolo et al., 2021; Stanojević et al., 2023). Neural signals at sentence-final positions could therefore reflect the increased syntactic integration associated with constituent completion, making bottom-up node count an important covariate when seeking to identify a neural signature of sentence-boundary detection.

Context, whether linear or syntactic, is not the only property that may systematically vary with sentence position. Lexical accessibility can also be influenced by context-independent properties of a word, most notably its frequency of occurrence. Word frequency has robust effects on language processing that are at least partly independent of contextual surprisal (Huizeling et al., 2022; Kretzschmar et al., 2015; Shain, 2024). Moreover, just as surprisal has been reported to decrease with sentence position, word frequency has been reported to increase with sentence position (Huizeling et al., 2022), such that sentence-final words may differ from other words in both contextual predictability and context-independent lexical accessibility.

Studying sentence processing in spontaneous speech presents a methodological challenge: unlike controlled experiments, naturalistic stimuli do not readily permit contrastive designs in which individual linguistic properties are manipulated while others are held constant. Here, we addressed this challenge using whole-brain parametric modeling to relate neural activity to multiple properties of the unfolding speech signal simultaneously. This approach served two complementary goals: to isolate responses associated with sentence boundaries beyond those accounted for by accompanying acoustic, prosodic, lexical, and syntactic properties, and to characterize neural sensitivity to these continuously varying properties during natural speech comprehension. To extend this second goal to semantic properties, we also included word concreteness. Although its relationship with sentence position is less established than that of lexical predictability or frequency, concreteness has been shown to modulate neural responses during semantic processing, including the N400, both in sentence contexts and during single-word processing (Barber et al., 2013; Holcomb et al., 1999; West & Holcomb, 2000). Inspired by EEG/MEG speech-tracking research that examine neural sensitivity to multiple levels of linguistic information (Gwilliams et al., 2025; Karunathilake et al., 2025; Ling et al., 2024), we pursued a related goal using a different method: whole-brain parametric modeling of BOLD responses to continuous speech properties. This approach allowed us to investigate sentence-boundary responses while also assessing the capacity of fMRI to reveal neural sensitivity to acoustic and linguistic variation during natural speech comprehension.

## 2. Methods

### 2.1. Participants

The present study used the publicly available *Merlin* and *Sherlock* subsets of the Narratives fMRI dataset (Nastase et al., 2021), which have previously been analyzed to address different research questions (Chang et al., 2022, 2024; Giglio et al., 2026; Zadbood et al., 2017). These subsets comprise functional MRI recordings from 36 adults aged 18–47 years (M = 21.7, SD = 4.7; 22 women) acquired while participants listened to naturalistic spoken narratives. The participants were right-handed native English speakers with normal or corrected-to-normal vision. All participants provided written informed consent in accordance with procedures approved by the Princeton University Institutional Review Board.

### 2.2. Speech Stimuli

The auditory stimuli consisted of spontaneous spoken recollections of episodes from the BBC television series *Merlin* and *Sherlock*. Both narratives were produced by the same female speaker, who watched each episode and subsequently recalled it aloud during fMRI scanning without prompts from the experimenter. The stimuli therefore comprised spontaneous speech rather than scripted or rehearsed narratives.

The *Merlin* stimulus file lasted 915 s in total and began with 25 s of music followed by 4 s of silence. The spoken story therefore lasted 886 s, corresponding to 591 functional volumes and included 2,245 transcribed words. The *Sherlock* stimulus file lasted 1081 s in total and likewise began with 25 s of music followed by 4 s of silence. The spoken story lasted 1,052 s, corresponding to 702 functional volumes and included 2,681 words. In both runs, the audio files were initiated 4.5 s after the onset of the functional scan. Together, the two spoken narratives provided 32 min 18 s of speech for analysis.

### 2.3. Experimental Procedure

All 36 participants listened to both spoken narratives. As part of the original experimental protocol (Zadbood et al., 2017), 18 participants viewed the *Merlin* episode and 18 viewed the *Sherlock* episode before the listening runs. Consequently, for each narrative, half of the participants had viewed the corresponding episode prior to the experimental session (the *movie* condition), whereas the other half had viewed the other episode and were therefore unfamiliar with the corresponding episode (the *naïve* condition). Following the scans, participants completed a free-recall task in which they were asked to describe events from the story in as much detail as possible. Recall quality was rated independently by three raters on a 1–10 scale, and the summed ratings were normalized by the maximum possible score to yield a comprehension score ranging from 0 to 1. Comprehension scores were available only for the *naïve* condition.

Participants listened through MRI-compatible insert earphones and were instructed to attend to the narratives while remaining still and keeping their eyes open. Presentation volume was adjusted as needed to ensure that participants could comfortably hear and understand the speech over the MRI acquisition noise.

### 2.4. MRI Acquisition

MRI data were acquired at the Princeton Neuroscience Institute using a 3-T Siemens Magnetom Skyra scanner equipped with a 20-channel phased-array head coil. Functional BOLD images were acquired using an interleaved T2*-weighted gradient-echo echo-planar imaging sequence with an in-plane GRAPPA acceleration factor of 2. Acquisition parameters were TR = 1,500 ms, TE = 28 ms, flip angle = 64°, bandwidth = 1,445 Hz/Px, in-plane resolution = 3 × 3 mm, slice thickness = 4 mm, matrix size = 64 × 64, and field of view = 192 × 192 mm. Twenty-seven axial slices were acquired without an interslice gap, providing approximately whole-brain coverage. Phase encoding was performed in the anterior-to-posterior direction. Three initial dummy volumes were acquired and discarded by the scanner to allow signal stabilization.

Anatomical images were acquired using a T1-weighted MPRAGE sequence with TR = 2,300 ms, TE = 3.08 ms, inversion time = 900 ms, flip angle = 9°, in-plane resolution = 0.859 × 0.859 mm, slice thickness = 0.9 mm, matrix size = 256 × 256, and 192 sagittal slices.

### 2.5. Data Annotation and Regressor Construction

Although the publicly available dataset included transcripts and word-level timestamps, we independently verified both to ensure accurate transcription and word timing. Two research assistants manually reviewed the transcripts and corrected transcription errors. The corrected transcripts were then force-aligned to the audio using the Montreal Forced Aligner (MFA; version 3.3.8; McAuliffe et al., 2017). A third research assistant inspected the alignments in Praat (Boersma et al., 2026), making further corrections to the transcripts and word timings. Across these stages, the final transcripts contained a net increase of 40 words relative to the original transcripts. The corrected timestamps were used to define a word-onset regressor and the timing of all lexical, prosodic, and syntactic predictors.

To isolate neural responses associated with sentence boundaries, we included a sentence-boundary regressor of interest alongside acoustic, lexical, semantic, prosodic, and syntactic regressors, as described below (Figure 1). The corrected and annotated transcripts, along with the annotated and calculated features, are available on Zenodo (Levy-David et al., 2026).

**Figure 1.**
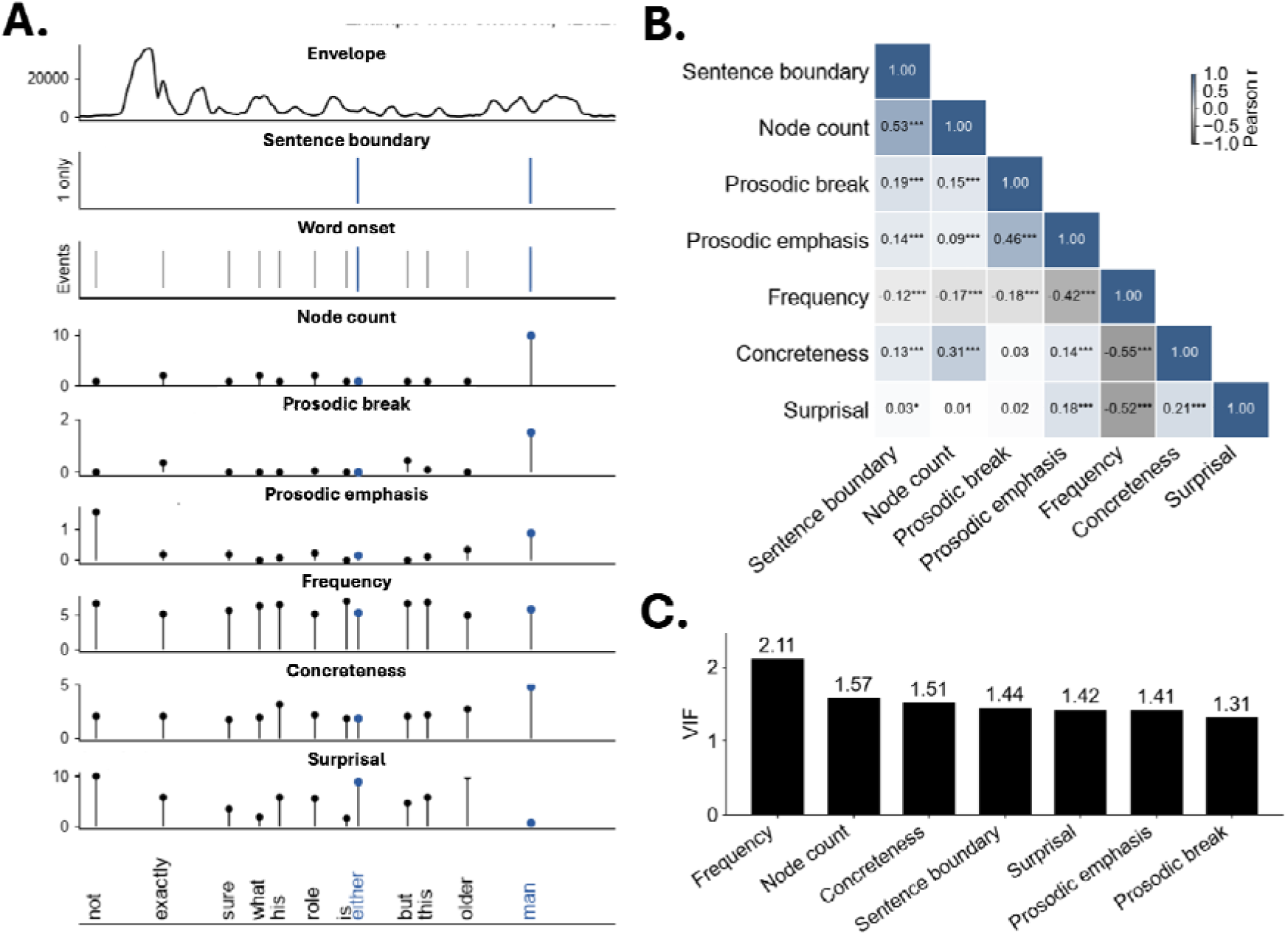
Relationships among sentence boundaries and linguistic properties. (A) Example segment from the *Merlin* spontaneous speech stimulus (465.5–470.5 s), illustrating the temporal alignment of the predictors included in the fMRI model. The corresponding transcript is shown along the bottom of the panel, allowing the word-level predictors to be viewed relative to the unfolding speech. Vertical lines indicate event onsets, and marker height reflects the value of each parametric predictor. Sentence-final words are highlighted in blue. (B) Pairwise Pearson correlations among sentence-boundary status, bottom-up node count, prosodic break, prosodic emphasis, lexical frequency, concreteness, and lexical surprisal. Asterisks indicate statistical significance (*p* < .05, \**p* < .01, \*\**p* < .001). (C) Variance inflation factors (VIFs) for the sentence-boundary and word-level parametric predictors included in the collinearity assessment.

#### Sentence Boundary Regressor

A sentence, intuitively understood as an independent entity conveying a complete idea, can be grammatically characterized as the maximal syntactic unit that does not depend on any other unit (Bloomfield, 1933, p. 170; Downing & Locke, 2006, p. 272). For example, “this is a sentence” is a complete, independent sentence, whereas “because this is a sentence” is not – the word “because” renders it a dependent clause that cannot stand alone. Following this definition, sentence boundaries were manually annotated by two trained research assistants based on the grammatical structure they perceived while listening to the recordings. A sentence boundary label was assigned to the word judged by the annotators as the final word of each complete sentence. Coordinated clauses joined by “and” or “but” were treated as separate sentences, provided that the coordination was not embedded within a larger clause. Final words of incomplete sentences were annotated separately for purposes of syntactic analysis (see *bottom-up node count* regressor), but were not considered sentence boundaries for the present regressor. Inter-rater agreement across the three categories (boundary, no-boundary, incomplete sentence) was 95.3% (Cohen’s κ = .76). When annotations were binarized as sentence boundary versus all other words, agreement was 96.5% (Cohen’s κ = .8). All disagreements were subsequently resolved through discussion to produce a consensus annotation. The final dataset consisted of 4,966 words (including filler words), of which 467 (9.4%) were sentence-final words; 98 words were final words of incomplete sentences and were not treated as sentence boundaries in the current analysis. Thus, the resulting sentence-boundary predictor consisted of events time-locked to the onset of each sentence-final word.

#### Word Onset Regressor

The word-onset predictor was binary, with events time-locked to the manually corrected onset of each word. This regressor modeled the general BOLD response associated with word occurrence, independently of other properties of individual words.

#### Acoustic Envelope Regressor

The acoustic envelope provides a continuous representation of the moment-to-moment amplitude structure of speech and differs from the remaining predictors in the model, which were primarily defined at discrete word onsets. Envelope tracking is widely used in EEG and MEG studies of continuous speech, where neural activity reliably follows fluctuations in the speech envelope (Aiken & Picton, 2008; Ding & Simon, 2012, 2013; Har-Shai Yahav & Zion Golumbic, 2021; Lalor & Foxe, 2010; Levy et al., 2025). Although the temporal resolution of fMRI is substantially lower, recent work has demonstrated that HRF-convolved speech envelopes can also predict BOLD activity in primary and non-primary auditory cortex (Hausfeld et al., 2024). Including this continuous acoustic regressor, therefore, allowed us to account for ongoing stimulus-driven fluctuations in the speech signal when estimating neural responses associated with sentence boundaries. A continuous broadband acoustic-envelope regressor was derived from each narrative using the filterbank-based Hilbert-envelope procedure employed in previous M\EEG studies of neural speech tracking (Crosse et al., 2016; Kaufman & Zion Golumbic, 2023). The audio waveform was filtered into 20 logarithmically spaced frequency bands extending from 100 Hz to 80% of the Nyquist frequency. Fourth-order Butterworth filters were applied bidirectionally to avoid phase distortion. Within each band, the magnitude of the analytic signal was calculated using the Hilbert transform, and the resulting narrowband envelopes were summed across bands. The broadband envelope was subsequently smoothed using a centered 20-ms moving-average window. The envelope was aligned to the fMRI acquisition timeline and adjusted to match the duration of the corresponding functional run. Before hemodynamic convolution, the aligned envelope was resampled to 100 Hz using polyphase resampling.

Unlike the other predictors, which were specified as event timings (sentence boundary, word onset) or event-linked amplitude modulators (see below), the acoustic envelope was represented as a continuously sampled signal. Continuously sampled signals have to be convolved with an HRF before supplying it to AFNI. Specifically, we convolved the envelope with a peak-normalized single-gamma hemodynamic response function kernel spanning 32 s, with a shape parameter of 6 and a scale parameter of 1. This parameterization was selected based on a preliminary cross-correlation analysis between the acoustic envelope and BOLD activity in primary auditory cortex. The convolved signal was averaged within successive non-overlapping 1.5-s windows corresponding to the fMRI TR, then demeaned and scaled to unit variance. The resulting time series was entered directly into the GLM without additional HRF convolution in AFNI.

#### Lexical Frequency Regressor

Lexical frequency was quantified using Zipf-scaled frequency estimates obtained with the *wordfreq* Python package [https://github.com/rspeer/wordfreq]. The package integrates frequency estimates derived from multiple large-scale text sources, including subtitles, encyclopaedic text, news, books, web text, and social media. Zipf frequency is defined as the base-10 logarithm of word frequency per billion words, *log_10_(p×10^9^)*, such that a one-unit increase corresponds to a tenfold increase in estimated word frequency. Higher values therefore indicate more frequent words. The resulting English Zipf values were assigned to the corresponding word forms and entered as a word-level parametric modulator.

#### Lexical Surprisal Regressor

Lexical surprisal was estimated using the GPT-2 autoregressive language model (Radford et al., 2019; 124M variant), implemented using the Hugging Face Transformers library. For each word except the first word in each file, surprisal was defined as the negative base-2 logarithm of its conditional probability given the preceding context, *-log_2_(p(w_i_|w_<i_)),* using up to GPT-2’s maximum context window of 1,024 tokens. Surprisal was calculated after removing annotated filler words (e.g., *uh*, *um*, *like*) and word repetitions from the input sequence. When a word was represented by multiple GPT-2 subword tokens, word-level surprisal was calculated as the sum of the surprisal values of its constituent tokens. The resulting values were assigned to the corresponding word forms and entered as a word-level parametric modulator.

#### Concreteness Regressor

Lexical concreteness was estimated using the *WordTangible* Python package [https://github.com/jrrobison1/wordtangible]. The package integrates normative concreteness ratings from the Brysbaert concreteness norms (Brysbaert et al., 2014), the Glasgow Norms (Scott et al., 2019), and the MRC Psycholinguistic Database (Coltheart, 1981). Ratings from these sources were averaged and normalized to a common 1– 5 scale, with higher values indicating greater concreteness. The resulting scores were assigned to the corresponding lexical items and entered as a word-level parametric modulator. Filler words and words for which no concreteness rating was available were excluded from this regressor.

#### Prosodic Break and Prosodic Emphasis Regressors

Two prosodic regressors––prosodic break and prosodic emphasis––were estimated using the Wavelet Prosody Toolkit (https://github.com/asuni/wavelet_prosody_toolkit; Suni et al., 2017). The toolkit combined contours of fundamental frequency, energy, and word duration using weights of 0.5, 0.5, and 1, respectively, and applied a continuous wavelet transform across multiple temporal scales. Prosodic break was then derived from the strength of linked wavelet minima, reflecting the prosodic break following each word. Emphasis, corresponding to the toolkit’s prominence measure, was derived from the strength of linked wavelet maxima, reflecting each word’s relative prosodic salience within the surrounding speech. Both measures were entered as continuous word-level parametric modulators.

#### Syntactic Node Count Regressor

Constituency parse trees were generated using the English constituency parser implemented in Stanza (version 1.11.1; Qi et al., 2020). The parser was given sentences based on the manual annotation (see Section *<u>Sentence Boundary</u> <u>Regressor</u>*), after removing filler words and repetitions, which were also annotated during the sentence segmentation process. The resulting trees were binarized using the Chomsky-normal-form transformation implemented in the Natural Language Toolkit (NLTK version 3.9.2; Bird et al., 2009). Syntactic node count was subsequently derived from each parse tree using in-house code. The code scored each word for the number of constituents whose right boundary coincided with that word. Under this bottom-up representation, higher node-count values indicated a greater number of syntactic constituents completed at that word and, consequently, greater syntactic integration load. When Stanza tokenized an original word into multiple tokens, node counts were summed across the corresponding tokens. Removed fillers and repetitions were assigned a node count of zero. The resulting values were aligned with the corrected word onsets and entered as a continuous parametric modulator.

To characterize the relationships among word-level speech properties, we calculated pairwise Pearson correlations (those involving binary sentence-boundary status were point-biserial correlations)^1^. We additionally fitted a binomial logistic regression with a logit link, using R’s *glm* function, predicting sentence boundary from lexical frequency (Zipf), surprisal, concreteness, prosodic emphasis, prosodic break, and bottom-up node count. Narrative was included as a fixed effect. Words with missing values for any predictor were excluded from the logistic regression, and all continuous predictors were standardized across the retained words. Model fit was assessed relative to a narrative-only model.

### 2.6. fMRI Analysis

Functional MRI data were preprocessed using AFNI (Cox, 1996; Cox & Hyde, 1997). The Merlin and Sherlock runs were processed separately using *afni_proc.py* (Reynolds et al., 2024). The first 22 volumes of each run (33 s), corresponding to the introductory music and subsequent silence, were excluded from the analysis. The retained time series began 0.5 s before speech onset, and all sentence- and word-level annotations were aligned to this time series.

#### Preprocessing

Preprocessing included slice-timing correction, alignment of the functional data to each participant’s T1-weighted anatomical image, spatial normalization to the TT_N27 Talairach template, motion correction, spatial smoothing with a 4-mm FWHM Gaussian kernel, masking, and signal scaling. Functional-to-anatomical alignment was performed using the *lpc+ZZ* cost function with the *giant_move* option. Motion correction was performed by registering volumes to the minimum-outlier volume, with alignment to the anatomical image incorporated into the transformation to standard space. The resulting functional data were masked using the anatomical mask. Volumes with motion exceeding 0.3 mm were censored, as were volumes in which more than 10% of voxels were identified as outliers, resulting in the exclusion of 5.5% (SD = 5.6%) of the functional data.

#### Subject-level analysis

The *Merlin* and *Sherlock* runs were analyzed separately for each participant using voxelwise general linear models implemented in AFNI’s *3dDeconvolve*, with all predictors included simultaneously (*Section 2.5*). Sentence boundaries constituted the primary regressor of interest and were modeled as events at the onset of sentence-final words. Word onsets were included as a control regressor. Both event regressors were convolved with AFNI’s canonical gamma (*GAM*) hemodynamic response function. Six additional word-level predictors—Zipf frequency, lexical surprisal, concreteness, prosodic break, prosodic emphasis, and bottom-up node count—were entered as parametric modulators, with each word’s predictor value modulating the amplitude of the modeled hemodynamic response time-locked to that word’s onset. These predictors were mean-centered prior to model estimation. The acoustic envelope was included as a continuous control regressor and was convolved with the hemodynamic response function prior to its inclusion in the GLM (see Section *<u>Acoustic Envelope Regressor</u>*). Six demeaned motion parameters and their temporal derivatives were included as nuisance regressors. Low-frequency signal drift was modeled using polynomials, with orders determined automatically based on run duration (sixth order for *Merlin* and eighth order for *Sherlock*).

#### Group-level analysis

Group-level effects were assessed using linear mixed-effects models implemented in AFNI’s *3dLMEr* (Chen et al., 2013, 2021). A separate model was fitted for each regressor, using two coefficient estimates per participant from the subject-level analyses: one from *Merlin* and one from *Sherlock*. Narrative-to-condition assignment was counterbalanced across participants, as described in *<u>Experimental Procedure</u>*. Each model used the subject-level beta estimates as the outcome, with condition (*naïve*/*movie*), comprehension score, age, and sex as fixed effects. A by-participant random intercept accounted for the repeated observations. Comprehension was represented by each participant’s score from the *naïve* condition, entered as a participant-level covariate. Comprehension scores and age were mean-centered. For each regressor, the group-level effect was taken as the average of the two estimated effects in the *movie* and *naïve* conditions.

Correction for multiple comparisons was performed using cluster-based inference with AFNI’s *3dClustSim* (Cox et al., 2017). AFNI’s *3dClustSim* provides a simulation-based cluster-level correction for multiple comparisons by generating a null distribution of the maximum cluster size under noise, with spatial autocorrelation properties estimated from the subject-level residuals. The spatial autocorrelation parameters were estimated separately from the residual time series of each participant using *3dFWHMx* with the *-acf* option and the participant-specific EPI-anatomical mask. The resulting autocorrelation parameters were averaged across participants and runs and entered into *3dClustSim* using the group-level mask. The resulting mean ACF parameters were 0.612, 3.115, and 9.175. With these parameters, *3dClustSim* estimated the number of contiguous voxels required at a given voxelwise uncorrected *p*-value to achieve a specified cluster-level corrected *p*-value. Using a two-sided test with a nearest-neighbor (NN1) clustering criterion, a voxelwise threshold of *p < .01* and a minimum cluster extent of 39 contiguous voxels yielded a cluster-level corrected significance threshold of *p < .05*.

## 3. Results

### 3.1. Relationships among sentence boundaries and linguistic predictors

As expected in natural speech, sentence boundaries co-occurred with variation in several linguistic and prosodic features. We therefore first examined the relationships among the predictors included in the fMRI model (Figure 1). Figure 1A illustrates an example segment of the stimulus, showing how sentence boundaries coincide with variation in syntactic structure, prosody, and lexical properties across the unfolding speech.

The strongest association with sentence boundaries was observed for bottom-up node count (*r* = .53; Figure 1B), consistent with the accumulation of constituent closures at sentence-final positions. Sentence boundaries were also positively associated with prosodic break (*r* = .19), concreteness (*r* = .13), and prosodic emphasis (*r* = .14), and negatively associated with lexical frequency (*r* = -.12). Their association with lexical surprisal was comparatively small (*r* = .03). Several predictors were also correlated with one another, most notably lexical frequency with concreteness (*r* = -.55) and surprisal (*r* = -.52), and prosodic break with prosodic emphasis (r = .46). To assess the predictors’ joint association with sentence boundaries, we fitted a logistic regression including all six word-level parametric predictors and narrative as a fixed effect. The analysis included 4,369 words after excluding 597 words with missing predictor values (460 disfluencies and 137 other words). The model showed a substantial improvement in fit over a narrative-only model (χ²(6) = 1,600.3, *p* < .001, McFadden’s pseudo-*R*² = .56). The only three properties that showed significant associations with sentence boundary were bottom-up node count (β = 1.97, SE = 0.09; *p* < .001; odds ratio = 7.2), word frequency (β = -.61, SE = 0.1; *p* < .001; odds ratio = .5) and prosodic break (β = .45, SE = 0.07; p < .001; odds ratio = 1.6).

Despite these associations, variance inflation factors were low across predictors, ranging from 1.31 to 2.11 (Figure 1C). The highest VIF was observed for lexical frequency (2.11), followed by bottom-up node count (1.57), concreteness (1.51), sentence boundary (1.44), surprisal (1.42), prosodic emphasis (1.41), and prosodic break (1.31), indicating limited multicollinearity among the predictors included in the model.

### 3.2 Neural responses to sentence boundaries and associated prosodic and syntactic features

We first focused on sentence-boundary responses and on the syntactic and prosodic predictors, which are the most directly related to sentence endings. Both syntactic node count and prosodic structure provide information that systematically accompanies sentence boundaries in continuous speech and may therefore contribute to neural responses observed at sentence-final positions. This relationship was also evident in the present stimuli, with bottom-up node count and prosodic break showing strong and significant associations with sentence boundaries (Figure 1B). We therefore examined sentence-boundary responses alongside these closely related features, while simultaneously accounting for the remaining acoustic, lexical and semantic properties included in the model.

**Sentence boundaries** were associated with a distributed pattern of significant BOLD responses (Figure 2A; Table 1). Negative boundary-related effects were predominantly left-lateralized, involving frontal and posterior temporoparietal regions, with the largest cluster centered in left frontal cortex (*k* = 627, *Z*max = -6.13). Positive effects showed a different spatial distribution and were particularly prominent in the right temporal lobe. The largest positive temporal cluster encompassed the right middle and superior temporal cortex (*k* = 261, *Z*max = 5.37). Smaller positive effects were also observed in right frontal and medial parts (see Table 1).

**Figure 2.**
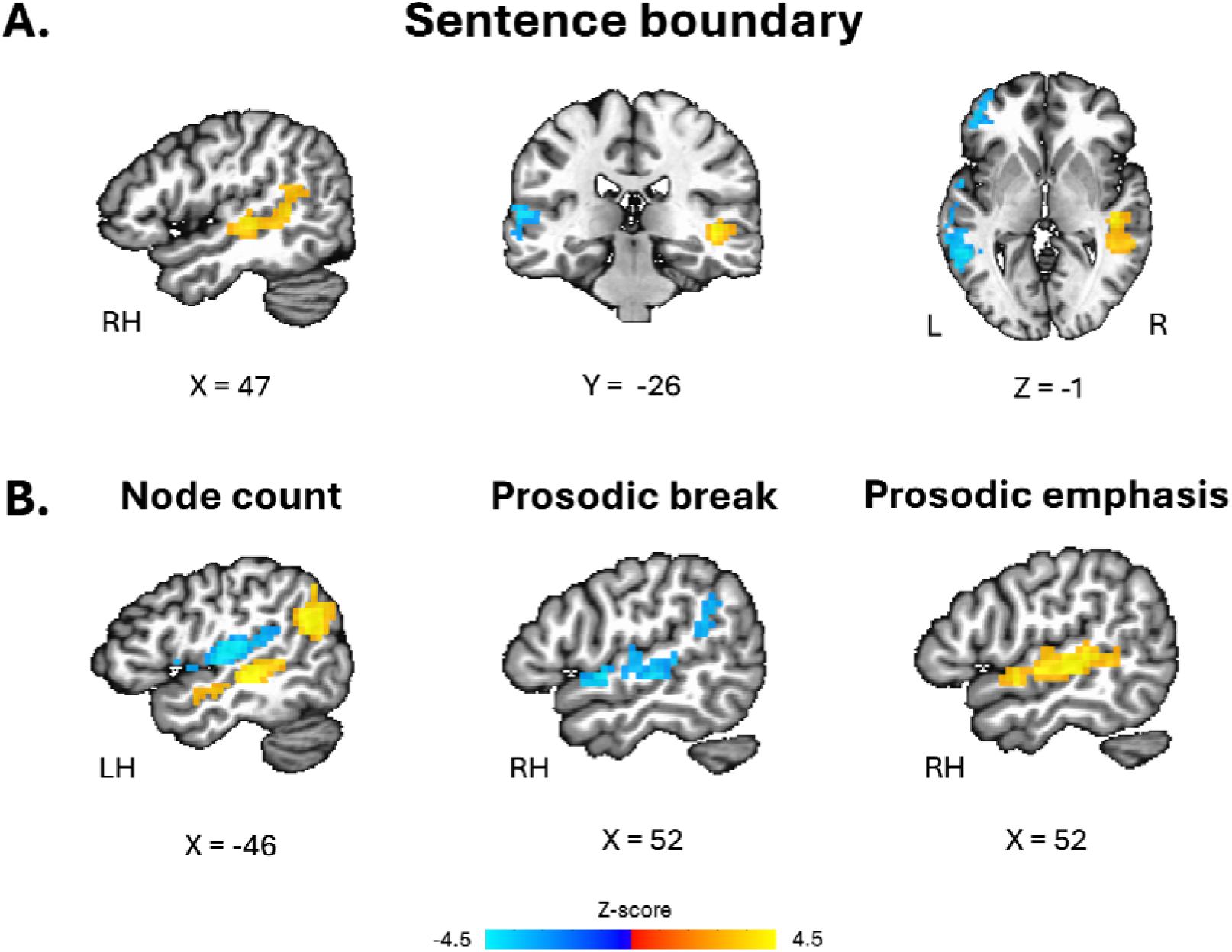
Neural responses to sentence boundaries and closely related syntactic and prosodic features. (A) Group-level response to sentence boundaries. (B) Group-level responses associated with bottom-up node count, prosodic break and prosodic emphasis. Warm colors indicate positive effects and cool colors indicate negative effects. Maps are shown as group-level *Z* scores, thresholded at voxelwise *p* < .01 and cluster-corrected *p* < .05 (NN1; minimum cluster extent = 39 voxels). Slice coordinates are shown in Talairach TT_N27 space. Full cluster-level results are reported in Table 1.

**Table 1.**
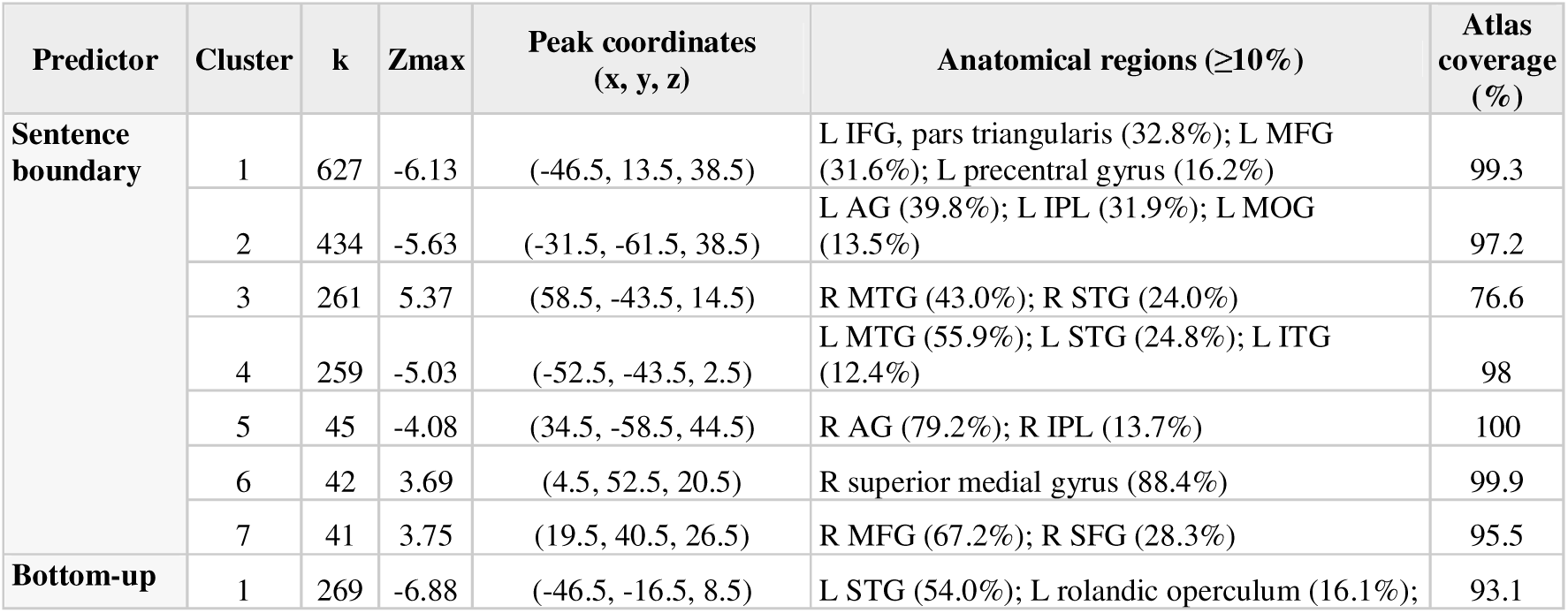
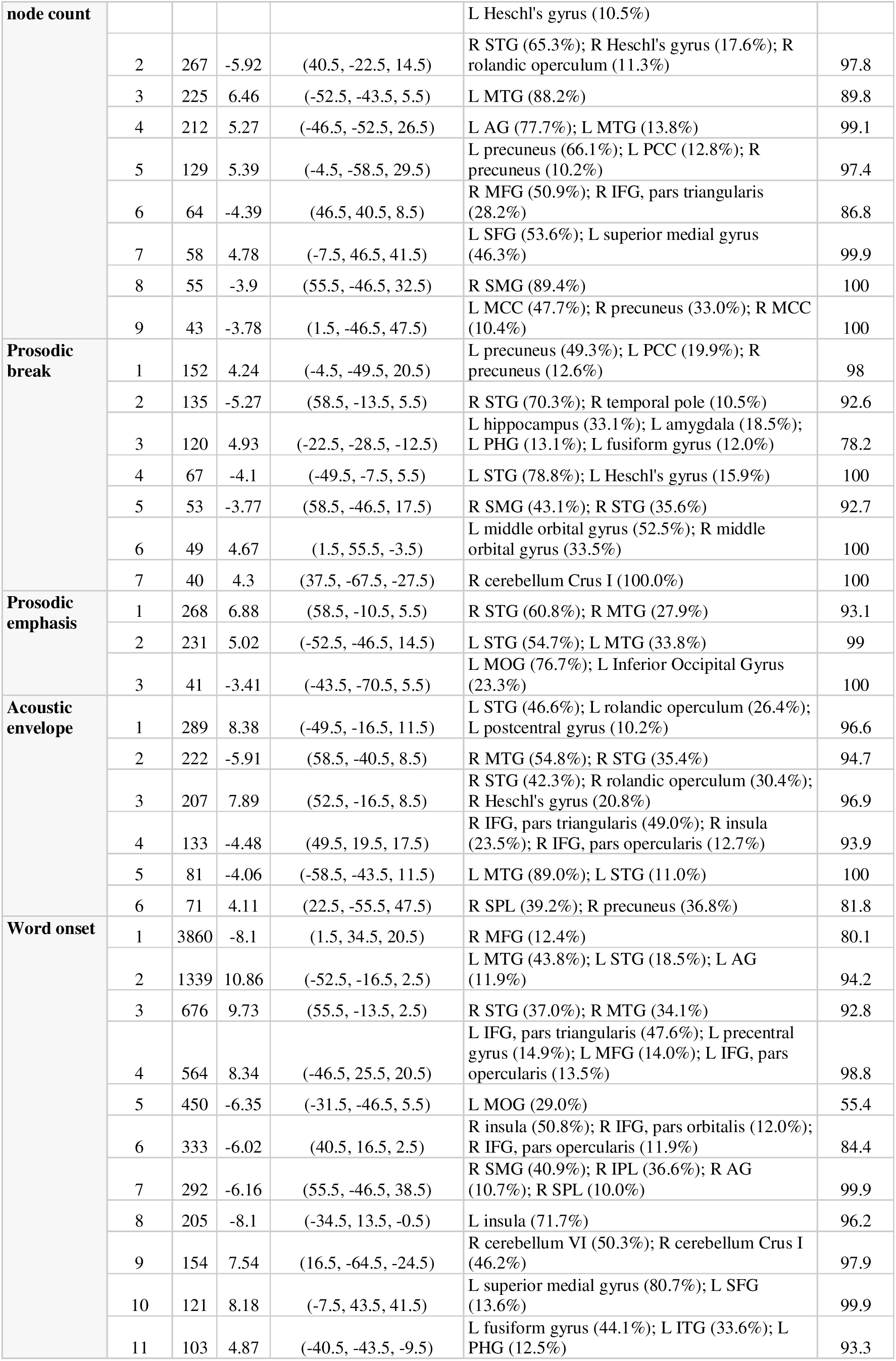
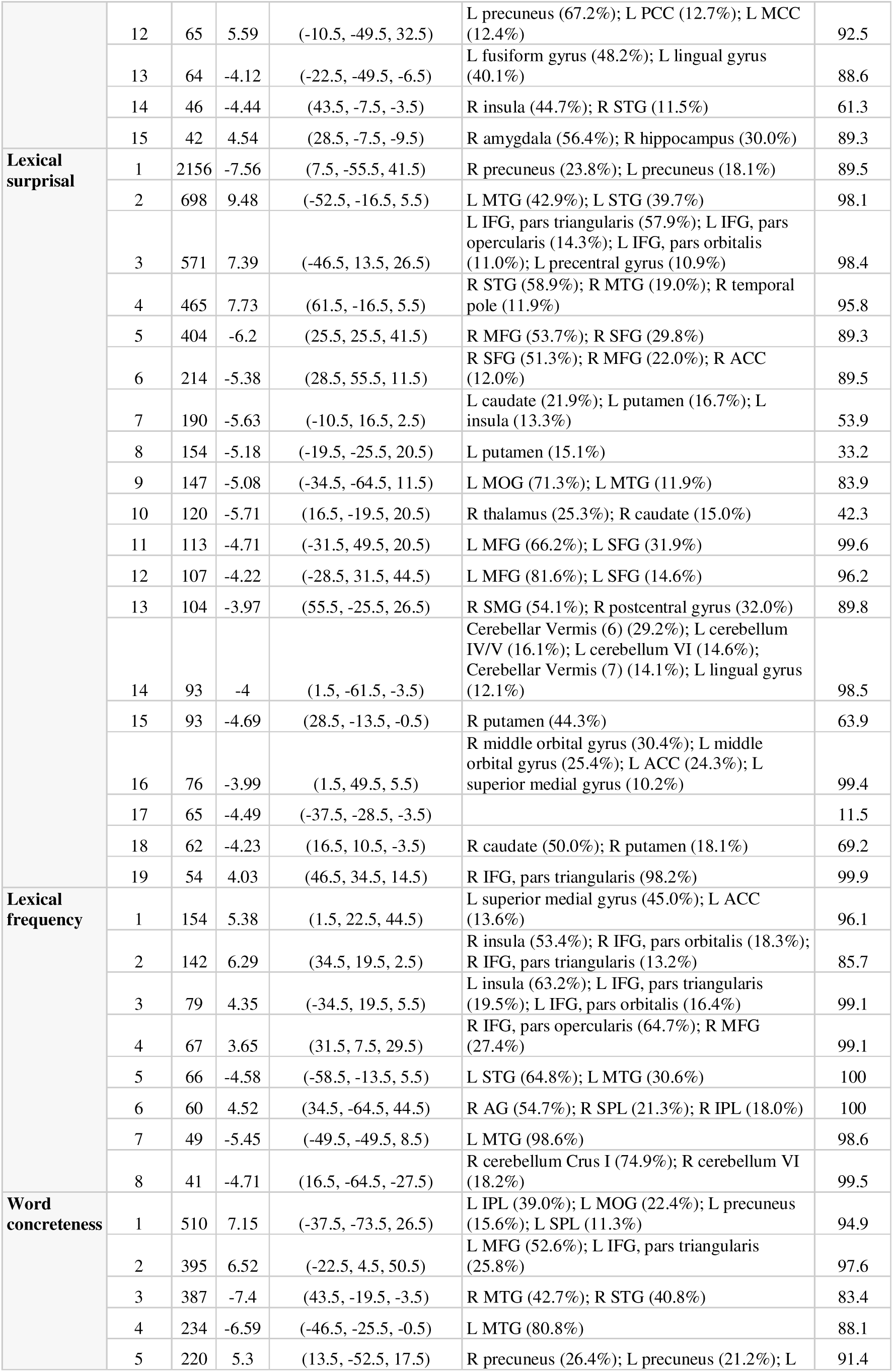
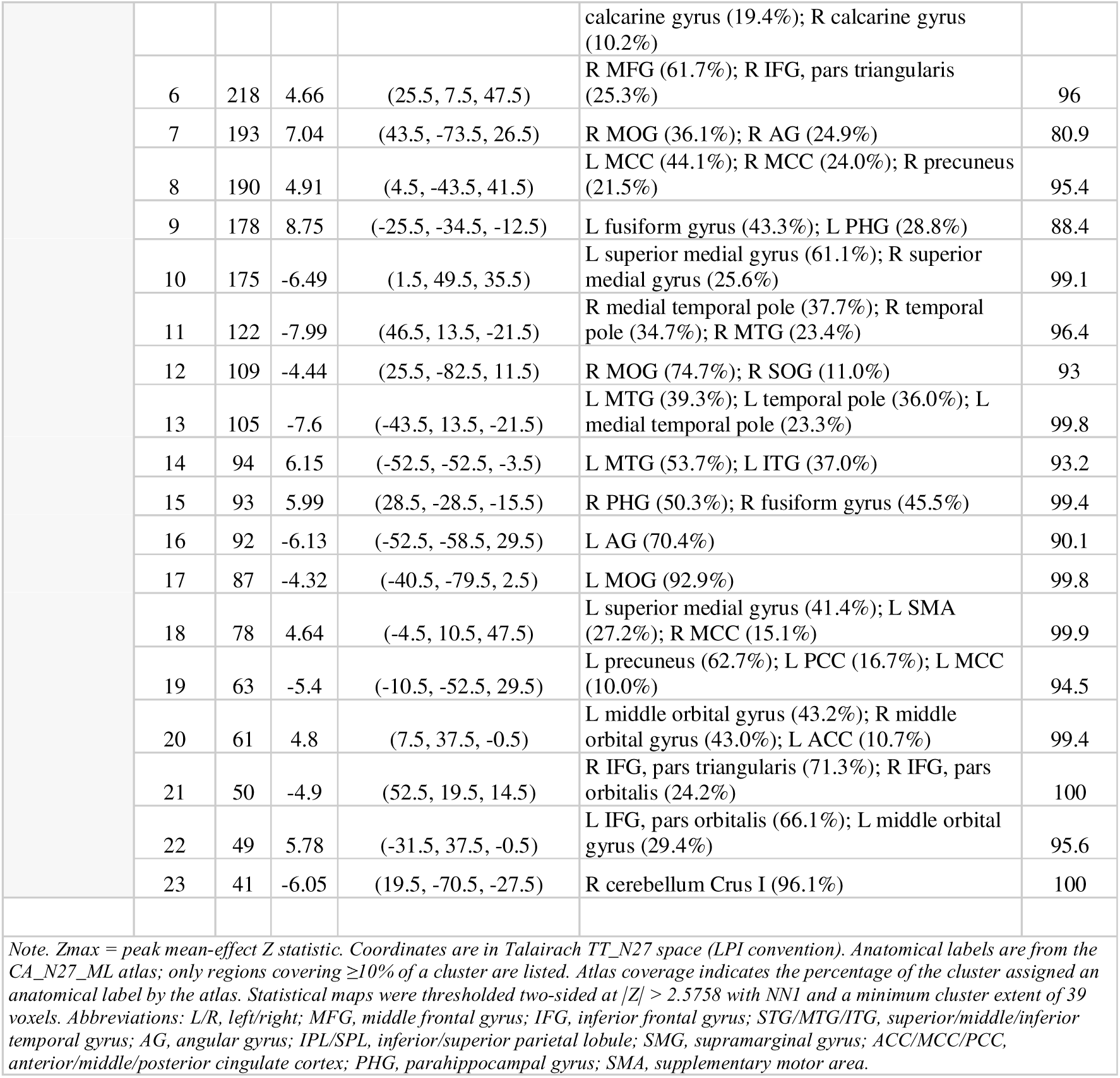
Whole-brain cluster results for all model predictors. Significant clusters are reported for the group-level mean effect of each predictor. For each cluster, the table shows cluster size (*k*), peak *Z* statistic, peak coordinates in Talairach TT_N27 space, and anatomical regions based on the CA_N27_ML atlas. The sign of the peak Z statistic indicates the direction of the effect. Only atlas regions accounting for ≥10% of the cluster are listed. Statistical maps were thresholded at voxelwise *p* < .01 and cluster-level corrected *p* < .05 (NN1; minimum cluster extent = 39 voxels).

The syntactic and prosodic predictors modeled alongside sentence boundaries also showed significant neural responses (Figure 2B; Table 1). **Bottom-up node count** was associated with predominantly left-lateralized positive responses, with the largest positive effects in the middle temporal gyrus (*k* = 225, *Z*max = 6.46) and the angular gyrus (*k* = 212, *Z*max = 5.27). These responses largely overlapped with regions showing negative associations with sentence boundary (Supplementary Figure S1A). Node count was also associated with prominent bilateral negative responses in superior temporal gyrus and Heschl’s gyrus, with the largest clusters in the left (*k* = 269, *Z*max = -6.88) and right (*k* = 267, *Z*max = -5.92). **Prosodic break** and **prosodic emphasis** likewise showed prominent temporal-lobe responses, but with different response profiles. In bilateral superior temporal cortex, prosodic break was associated predominantly with negative BOLD responses, strongest and largest in the right STG (*k* = 135, *Z*max = -5.27), whereas prosodic emphasis was associated with strong positive responses bilaterally, with its largest effect in the right STG/MTG (*k* = 268, *Z*max = 6.88). Responses associated with the two prosodic regressors in the right temporal lobe did not spatially overlap with regions showing positive sentence-boundary effects (Supplementary Figures S1B and S1C).

### 3.3 Neural responses to acoustic and lexical properties of speech

We next examined the remaining predictors in the model, which captured more general acoustic and lexical properties of the unfolding speech signal (Figure 3; Table 1). These predictors showed robust and spatially differentiated neural responses across multiple levels of speech processing.

**Figure 3.**
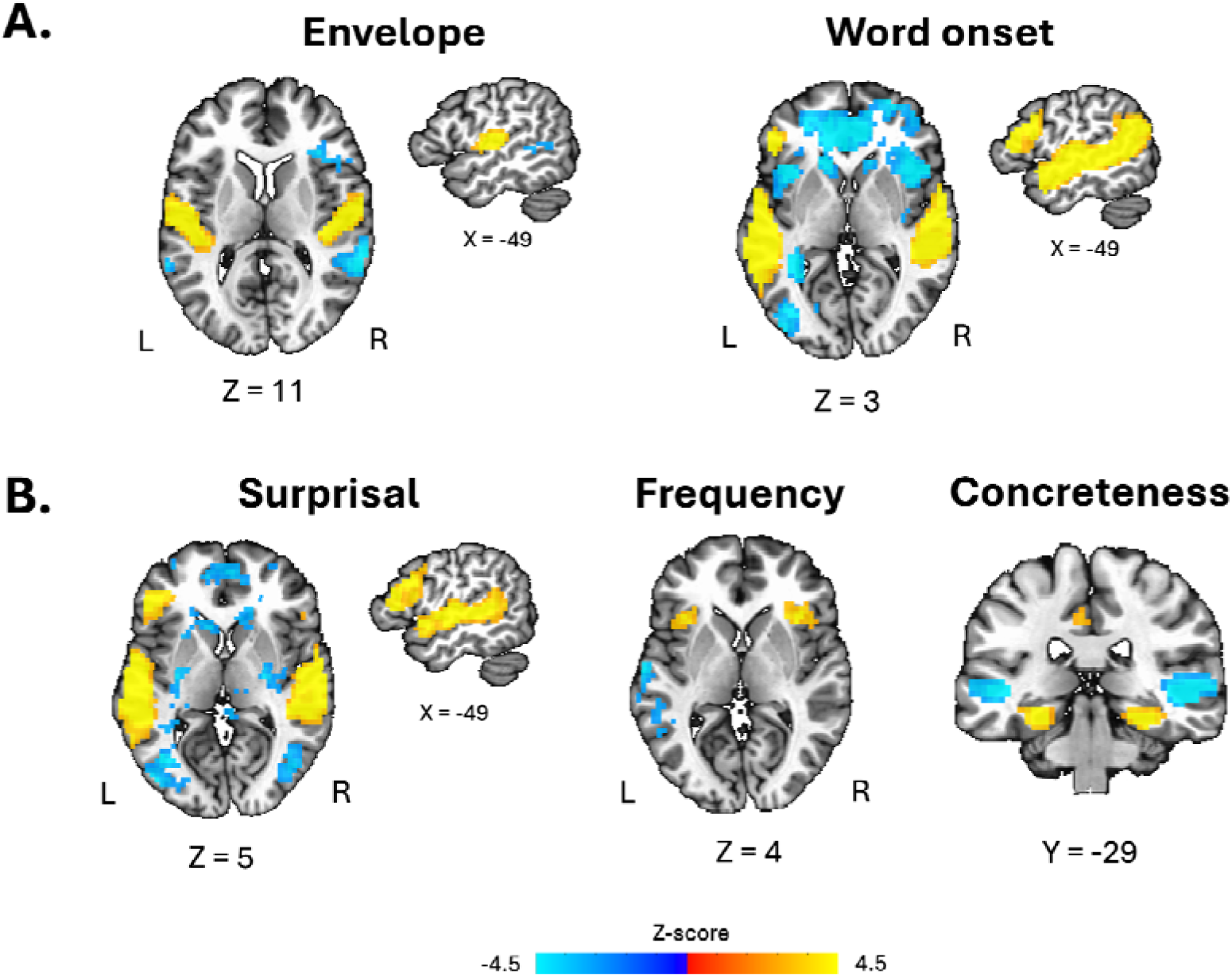
Neural responses associated with acoustic and lexical properties of spontaneous speech. **(A)** Group-level responses associated with the acoustic envelope and word onset. **(B)** Group-level responses associated with lexical surprisal, lexical frequency, and concreteness. Warm colors indicate positive effects and cool colors indicate negative effects. Maps are shown as group-level *Z* scores, thresholded at voxelwise *p* < .01 and cluster-corrected *p* < .05 (NN1; minimum cluster extent = 39 voxels). Slice coordinates are shown in Talairach TT_N27 space. Full cluster-level results are reported in Table 1.

The **acoustic envelope** was associated most prominently with positive bilateral responses in superior temporal and perisylvian auditory regions (Figure 3A). The strongest effects were observed in the left superior temporal cortex (k = 289, Zmax = 8.38) and in a corresponding right-hemisphere cluster (k = 222, Zmax = -5.91) extending across the superior temporal cortex and Heschl’s gyrus (k = 207, Zmax = 7.89).

**Word onsets** elicited a considerably widespread pattern of responses, including strong positive effects throughout bilateral temporal cortex, extending into left inferior frontal regions, together with additional frontal and medial responses. Negative effects were distributed across parietal, occipital, and insular regions (Figure 3A; Table 1). The largest positive effects were observed in left temporal cortex (k = 1,339, Zmax = 10.86) and right superior and middle temporal cortex (k = 676, Zmax = 9.73).

Among the lexical predictors (Figure 3B), **surprisal** produced the most extensive response pattern. Higher surprisal was associated with strong positive responses in bilateral temporal language regions, with the largest left-hemisphere clusters in the temporal cortex (k = 698, Zmax = 9.48) and the inferior frontal gyrus (k = 571, Zmax = 7.39), and a corresponding right temporal cluster (k = 465, Zmax = 7.73). Widespread negative effects were observed in precuneus, in frontal regions and in the basal ganglia. Lexical **frequency** showed a more spatially restricted pattern, with positive effects primarily in the insula both in the right (k = 142, Zmax = 6.29) and left (k = 79, Zmax = 4.35) hemispheres, and negative responses in the left superior temporal gyrus (k = 66, Zmax = -4.58). **Concreteness** was associated with a broad pattern of positive and negative responses across temporal, frontal, parietal, and ventral temporal regions (Table 1), with notable negative activations (i.e., stronger responses to abstract words) in bilateral mid-superior temporal gyri, and positive activations (i.e., stronger responses to concrete words) in bilateral parahippocampal gyri (Figure 3B).

## 4. Discussion

The present study asked whether the brain identifies sentences in continuous speech. Beyond its relevance to the neural mechanisms of comprehension, this question bears directly on the psychological status of sentences, which occupy a central place in linguistic theory. Although some previous neuroimaging studies have included sentence-boundary regressors as covariates (e.g., Giglio et al., 2024; Shain et al., 2022), we placed sentence-boundary detection at the center of our investigation. Specifically, we asked whether sentence endings evoke a neural response beyond that accounted for by accompanying acoustic, prosodic, lexical, semantic and syntactic properties. We found such a response in both left and right temporal regions, consistent with the possibility that the brain actively marks sentence completion during continuous speech comprehension.

Negative association between **sentence-final words** and BOLD responses were found in left posterior parts of the middle temporal gyrus and in the left angular gyrus. This negative association may reflect reduced engagement of structure-building processes once a sentence is complete. These regions also showed positive associations with bottom-up node count (Supplementary Figure 1A), consistent with their involvement in combining words into larger syntactic units. One possible interpretation is that completing a sentence reduces the need to maintain and extend its syntactic structure, allowing these processes to disengage. Because bottom-up node count was included in the same model, the negative sentence-final effect may reflect this reduction beyond the structure-building demands associated with closing constituents at the final word.

The positive sentence-boundary response in the right hemisphere, however, was unexpected. We had anticipated positive boundary-related responses primarily in the left hemisphere, given its established involvement in the processing of language and linguistic structure (Friederici, 2011; Geschwind, 1970; Malik-Moraleda et al., 2022). Consistent with that expectation, Giglio et al. (2024), who analyzed the same dataset (but for different purposes), did report positive sentence-boundary effects in the left inferior frontal cortex (BA45) and posterior middle temporal cortex. Our negative left-hemisphere effects therefore stand in direct contrast with their findings, while our whole-brain analysis additionally identified positive responses in the right hemisphere.

Several methodological differences may help explain the opposite effects in left temporal cortex and the additional right-hemisphere response identified here. First, Giglio et al. (2024) restricted their analysis to left-hemisphere regions of interest, whereas our whole-brain analysis allowed us to detect responses in either hemisphere. Second, our analysis was specifically designed to assess boundary-related responses. Thus, while Giglio et al. (2024) accounted for word frequency, surprisal, and bottom-up node count, we included additional relevant covariates, such as prosodic features. Finally, the studies differed in their operational definition of sentence boundaries. Giglio et al. (2024) treated any interruption of syntactic structure (i.e., restarts, parenthetical insertions, and repetitions) as points at which a new sentence began. Our sentence-boundary regressor, by contrast, marked only the final words of grammatically complete sentences; endings of incomplete sentences were annotated separately and excluded from this regressor. Their effects may therefore reflect a broader range of structural interruptions than those captured by our boundary definition. We consider possible functional interpretations of the right-hemisphere response in *Section 4.2*.

### 4.1 Tracking multiple levels of speech with fMRI

Before interpreting the right temporal response to sentence boundaries, we highlight the broader methodological contribution of this study: using fMRI to characterize neural sensitivity to multiple properties of continuous speech within a single model. We combined a continuously sampled acoustic-envelope regressor with event-linked parametric modulators representing prosodic, lexical, semantic, and syntactic properties (see also Brennan et al., 2016; Giglio et al., 2024, 2026 for similar approaches). Although these properties vary rapidly and often covary in naturalistic speech, modeling them jointly revealed their associations with BOLD activity while accounting for one another. These findings highlight a capability of fMRI that its limited temporal resolution can obscure: sensitivity to speech properties that fluctuate much faster than the hemodynamic response itself. Below, we discuss how acoustic and linguistic variation on millisecond-to-second timescales is reflected in the slower BOLD signal, allowing us to map where these properties are represented during natural speech comprehension.

One of the most striking findings was the robust association between the **acoustic envelope** and BOLD responses in the bilateral primary auditory cortices. This result is particularly informative given the marked difference in temporal scale between the acoustic signal, which fluctuates continuously on a millisecond timescale, and the hemodynamic response, which integrates neural activity over seconds. Nevertheless, temporal variation in the envelope explained reliable variance in auditory-cortical activity, even at the individual-participant level (see Supplementary Figure S2). This spatial pattern is consistent with extensive EEG and MEG work showing that neural activity in auditory cortex tracks the temporal envelope of continuous speech (Aiken & Picton, 2008; Ding & Simon, 2012; Kaufman & Zion Golumbic, 2023; Lalor & Foxe, 2010; O’Sullivan et al., 2015; Zion Golumbic et al., 2013). More recently, Hausfeld et al. (2024) demonstrated that envelope-related dynamics can also be recovered with fMRI, reporting speech-envelope tracking in bilateral Heschl’s gyrus as well as non-primary auditory regions. Our findings converge with this spatial pattern and extend this approach to a multivariate model of continuous speech, in which the contribution of acoustic information was estimated while simultaneously accounting for multiple linguistic properties of the speech. Some fMRI studies also incorporated closely related acoustic-energy measures, such as sound power, but they did that primarily as control predictor within an ROI analysis, without discussing their associated effects (Brennan et al., 2016; Shain et al., 2020, 2022), limiting direct comparison with the whole-brain effects observed here. The present findings further show that envelope-related BOLD variation can be detected in the expected auditory regions during ongoing speech, highlighting the potential of this measure for linking electrophysiological and hemodynamic approaches to speech tracking.

The lexical predictors of **word onset** and **lexical surprisal** were associated with widespread responses across the language network, consistent with previous findings (Brennan et al., 2016; Giglio et al., 2026; Russo et al., 2020; Shain et al., 2020). The **word frequency** effect is harder to situate within the existing literature, which reports heterogeneous neural correlates (Giglio et al., 2026; Mei et al., 2025; Sánchez et al., 2023; Wilson et al., 2009). Notably, Giglio et al. (2026), who analyzed the same dataset, reported small positive frequency-related clusters in left temporal and frontal cortices. Our analysis, however, points to positive clusters in the insulae and negative clusters in left temporal regions. Differences in such small clusters may reflect differences in model specification and statistical analysis. To note, we also found lexical frequency to be negatively associated with sentence boundaries, opposite to a previous report on an increase of word frequency with sentence position (Huizeling et al., 2022).

The semantic predictor of **concreteness** showed a topographic correspondence with the dorsolateral-to-ventromedial organization: lower concreteness was associated with stronger responses in lateral and superior temporal regions, whereas increasing concreteness was associated with stronger responses in ventral and medial temporal regions, including fusiform and parahippocampal cortex. This pattern resembles the organization of abstract and concrete words along the mid-temporal cortex reported by Hoffman et al. (2015), consistent with meta-analytic evidence demonstrating a preferential role of the parahippocampal gyrus in the representation of concrete words (Wang et al., 2010). Importantly, whereas Hoffman et al. (2015) demonstrated this organization using categorical contrasts in a controlled semantic task, the present study recovered a similar spatial organization using a continuous concreteness regressor during naturalistic speech comprehension. This convergence demonstrates that concreteness-related neural organization can be studied through graded variation in ongoing speech, providing a framework for studying lexical-semantic processing under more naturalistic conditions. It also suggests that the brain remains sensitive to the concreteness of individual concepts as they occur within a meaningful linguistic context, even when listeners are not explicitly asked to evaluate their semantic properties.

Prosodic emphasis, prosodic breaks, and bottom-up node count were particularly relevant covariates because sentence boundaries are associated with both prosodic changes and syntactic constituent completion (Brennan et al., 2016; Buxó-Lugo & Watson, 2016; Cooper & Paccia-Cooper, 1980; Giglio et al., 2024). Our analysis identified neural responses associated with these predictors in expected regions. **Bottom-up node count**, which indexes constituent completion and serves as a measure of syntactic integration demands, was associated with positive responses in left temporal and temporoparietal regions, including the middle temporal gyrus, and the angular gyrus, in line with previous research (Brennan et al., 2016; Giglio et al., 2024; Lopopolo et al., 2021; Stanojević et al., 2023). This localization contrasts with the right temporal response associated with sentence boundaries, suggesting that the two predictors capture different aspects of sentence processing. Noteworthy, this effect was found after accounting for LLM-derived surprisal, which highlights the relevance of explicitly hierarchical measures of syntactic structure beyond word-level predictability in explaining neural responses during language comprehension.

The two **prosodic predictors**, **emphasis** and **break**, captured changes in prosody as speech unfolded and were computed automatically from a combination of pitch, amplitude, and duration features. Both were associated with BOLD responses in the right superior temporal gyrus, with a positive association for emphasis and a negative association for break. Although interpreting these opposite directions requires caution, the localization of both effects is consistent with evidence implicating right temporal regions in the processing of speech prosody, particularly slow prosodic modulations (Meyer et al., 2002).

These findings provide a context for returning to our central research question: the sentence-boundary response in the right superior temporal gyrus.

### 4.2 The response of the right temporal cortex to sentence boundaries

Although we initially expected the sentence-boundary predictor to be associated with positive left-hemisphere responses (Giglio et al., 2024), the right temporal response suggests at least two possible interpretations. First, given the topography of the prosodic effects reported above, this right-hemispheric sentence-boundary response may reflect sensitivity to prosodic properties that covary with sentence boundaries but were not fully captured by our prosodic predictors. Second, it may reflect perceived prosody that is triggered by the syntactic structure. Third, persistence of the response in the right hemisphere after accounting for the modeled speech properties may reflect a contribution of right temporal regions to identifying sentence boundaries or marking the completion of sentence-level units. These interpretations are not mutually exclusive, and we consider each below.

The most immediate possibility is that our prosodic regressors did not capture all relevant prosodic variation. Because the sentence-boundary regressor was defined grammatically, any residual prosodic contribution would concern unmeasured properties systematically associated with grammatical sentence endings. Although our measures of prosodic break and emphasis summarize important aspects of linguistic prosody, they may not fully represent the combinations of pitch movement, temporal organization, and intensity through which speakers may also signal sentence completion. Future work could therefore compare alternative prosodic representations, incorporating explicit measures of pitch contours, preboundary lengthening, pauses, and pitch reset, together with their interactions and variation over longer stretches of speech. A complementary approach would be to incorporate ToBI-based prosodic annotations, which distinguish intonational categories such as pitch accents, phrase accents, boundary tones and hierarchical prosodic phrasing, and can be derived automatically (Beckman & Ayers Elam, 1997; Zhai & Hasegawa-Johnson, 2023). These annotations could help determine whether sentence-boundary responses reflect particular intonational configurations beyond the boundary-strength and emphasis measures used here. Ongoing work by one of the co-authors (GA) likewise aims to improve prosodic models for sentence-boundary detection. Testing whether the neural sentence-boundary response persists across these alternative representations would help clarify the contribution of prosodic information to the measured right hemispheric response to grammatical sentence boundaries. Importantly, a residual prosodic contribution would not make the effect unrelated to sentence structure, but it would limit our ability to attribute it specifically to boundary detection.

A separate but compatible possibility is that the right-hemisphere response reflects prosodic *expectations* shaped by linguistic structure, rather than solely the prosodic cues present in the acoustic signal. Prosodic breaks and syntactic boundaries do not align consistently in spontaneous speech (Bannon et al., 2026; Blaauw, 1994; Face, 2003), and listeners may use grammatical information to infer prosodic grouping when acoustic cues are weak or absent (Buxó-Lugo & Watson, 2016; Itzhak et al., 2010). Our sentence-boundary effect could therefore reflect a prosodic response shaped by the grammatical structure of the unfolding speech, which would not necessarily be captured by predictors derived from the acoustic signal alone.

A third interpretation is that the right temporal response reflects the integration of information into larger perceptual or linguistic units. This interpretation draws on evidence for right-hemisphere contributions to processing broader temporal patterns and global structure across several domains. In audition, the asymmetric sampling in time hypothesis proposes relatively longer temporal integration windows in right than left auditory regions, consistent with greater sensitivity to slow temporal modulations in the right hemisphere (Boemio et al., 2005; Oderbolz et al., 2025; Poeppel, 2003). In vision, a related hemispheric asymmetry has been reported, with a right-hemisphere advantage for processing global rather than local stimulus features (Fink et al., 1996; Robertson et al., 1988). In language, right-hemisphere contributions have been associated with distant semantic relationships (Beeman et al., 1994) and the use of context and macrostructure to construct coherent discourse interpretation (Johns et al., 2008; Lindell, 2006; Vigneau et al., 2011). Although these findings concern different representational levels and timescales, they motivate the possibility that right temporal regions contribute to recognizing when an unfolding sequence constitutes a complete linguistic unit. On this account, the sentence-boundary response could reflect the convergence of acoustic, prosodic, lexical, semantic and syntactic information signaling sentence completion. Such a representation could, in turn, provide feedback to other language regions, helping organize the ongoing speech stream into sentence-level units. These mechanisms remain speculative: the present findings identify a boundary-associated response but do not establish a mechanistic account. Future work could test how right temporal regions interact with the broader language network as evidence for sentence completion accumulates.

To conclude, the present study identified a response to sentence-final words in the right superior temporal gyrus beyond that explained by measured acoustic, prosodic, lexical, semantic and syntactic properties. More broadly, our findings demonstrate how fMRI can reveal neural sensitivity to multiple levels of continuous speech within a single model. By making sentence boundaries the focus of this analysis, we identified a response consistent with the brain treating sentences as cognitive units and actively marking their completion. Future research should test whether this response persists with more comprehensive prosodic models and distinguish the contributions of acoustic cues and linguistically driven prosodic expectations. Such work will help clarify how listeners identify sentence-level units within the unfolding speech signal.

## Declaration of generative AI and AI-assisted technologies in the manuscript preparation process

During the preparation of this work, the authors used ChatGPT Plus to assist with aspects of the data analysis and language editing. The authors reviewed and edited the output as needed and take full responsibility for the content of the published article.

## Supplementary

**Figure S1.**
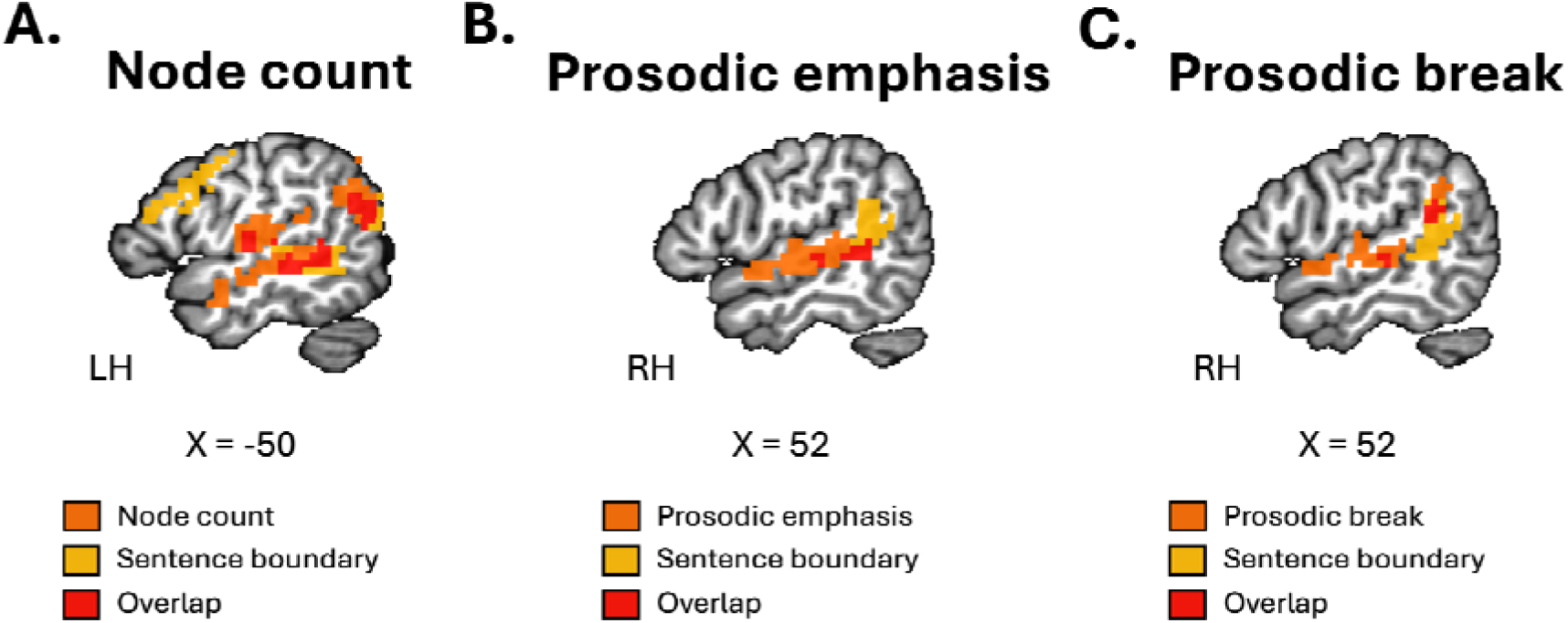
Spatial overlap between sentence-boundary responses and syntactic and prosodic predictors. **(A)** Overlap between sentence-boundary and bottom-up node count responses. **(B)** Overlap between sentence-boundary and prosodic emphasis responses. **(C)** Overlap between sentence-boundary and prosodic break responses. Orange indicates significant responses associated with the respective predictor, yellow indicates significant sentence-boundary responses, and red indicates spatial overlap between the two thresholded maps. Cluster masks were extracted after thresholding statistical maps at voxelwise *p* < .01 and cluster-corrected at NN1 with a minimum cluster extent of 39 voxels. Slice coordinates are shown in Talairach TT_N27 space.

**Figure S2.**
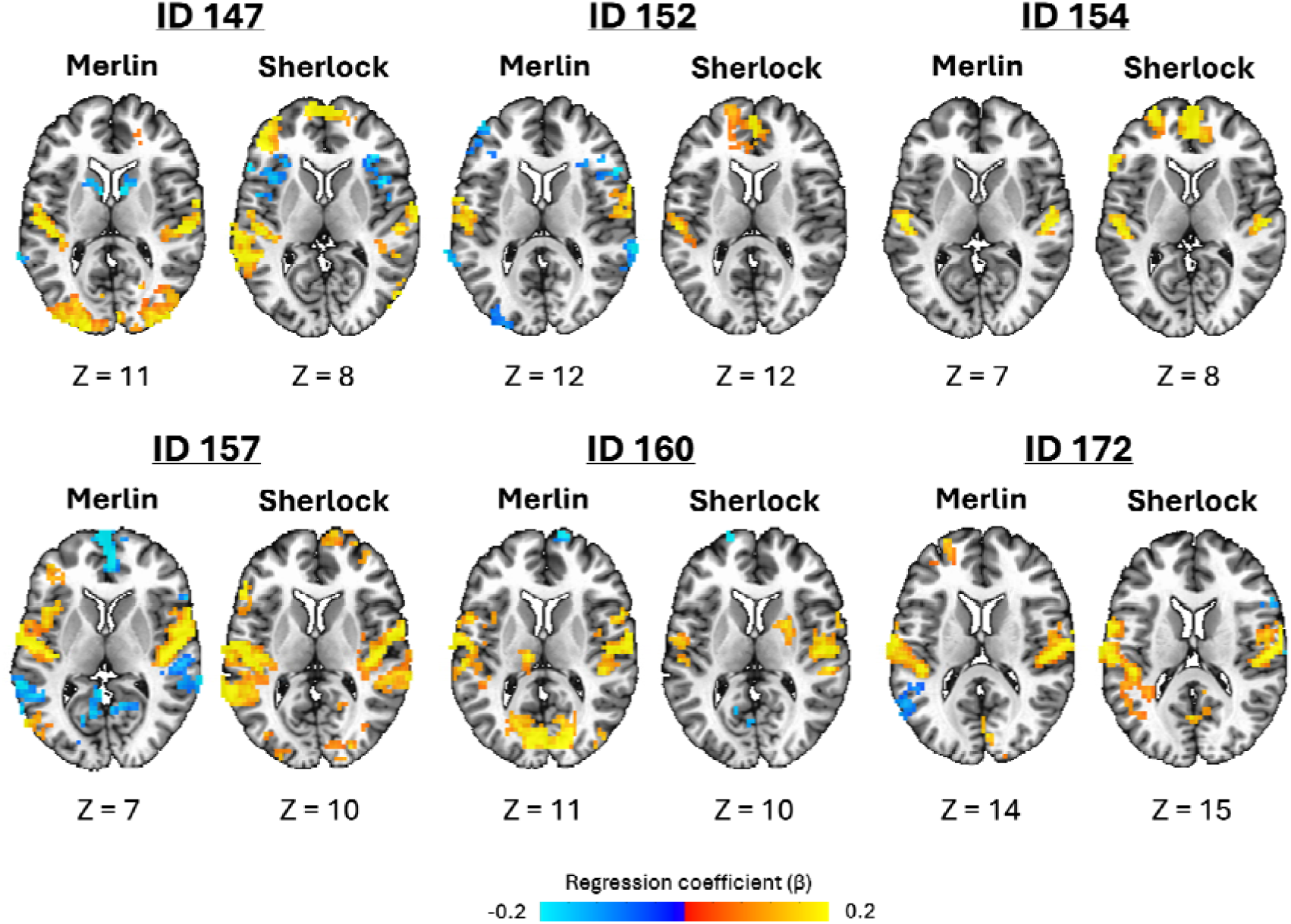
Individual-participant responses associated with the acoustic envelope across narratives. Statistical maps are shown for six representative participants for the *Merlin* and *Sherlock* narratives. Warm colors indicate positive effects and cool colors indicate negative effects. Maps are shown as individual-participant regression coefficients (β), thresholded at voxelwise *p* < .01 and cluster-corrected at NN1 with a minimum cluster extent of 39 voxels. Only effects with |β| ≥ 0.2 are displayed. Slice coordinates are shown in Talairach TT_N27 space.

## Footnotes

1 These correlations describe the speech properties themselves, before HRF convolution, rather than the columns of the fMRI design matrix. Therefore, word onset or the continuously sampled acoustic envelope were not included in this word-level correlation analysis.

## Notes

This research was supported by the Israel Science Foundation (543/25, recipient: G.A.).

https://zenodo.org/records/23342920?preview=1&token=eyJhbGciOiJIUzUxMiJ9.eyJpZCI6IjQyMzBmMDNiLWUwZjEtNDhmNC1hMzA0LWEwYzdkN2M1Y2NhYSIsImRhdGEiOnt9LCJyYW5kb20iOiI5YTM3ZWU2OTI0NWU2MGQ2MDAyYzc3NDQ2NzQ5ZTMxYiJ9.dzsygx1MS2ogIb9Qzxqqzc4htJHZJ2XjL1IUkRsSdXYy9h7xIrrNZGeyF7lK5lMX_ke-KEbbQKcMTcgxbRGLCg

### Competing Interest Statement

The authors have declared no competing interest.

