## Supplemental Figures for "Speech Tracking in fMRI Reveals a Distinct Response to Sentence Boundaries During Comprehension"

**Supplementary**

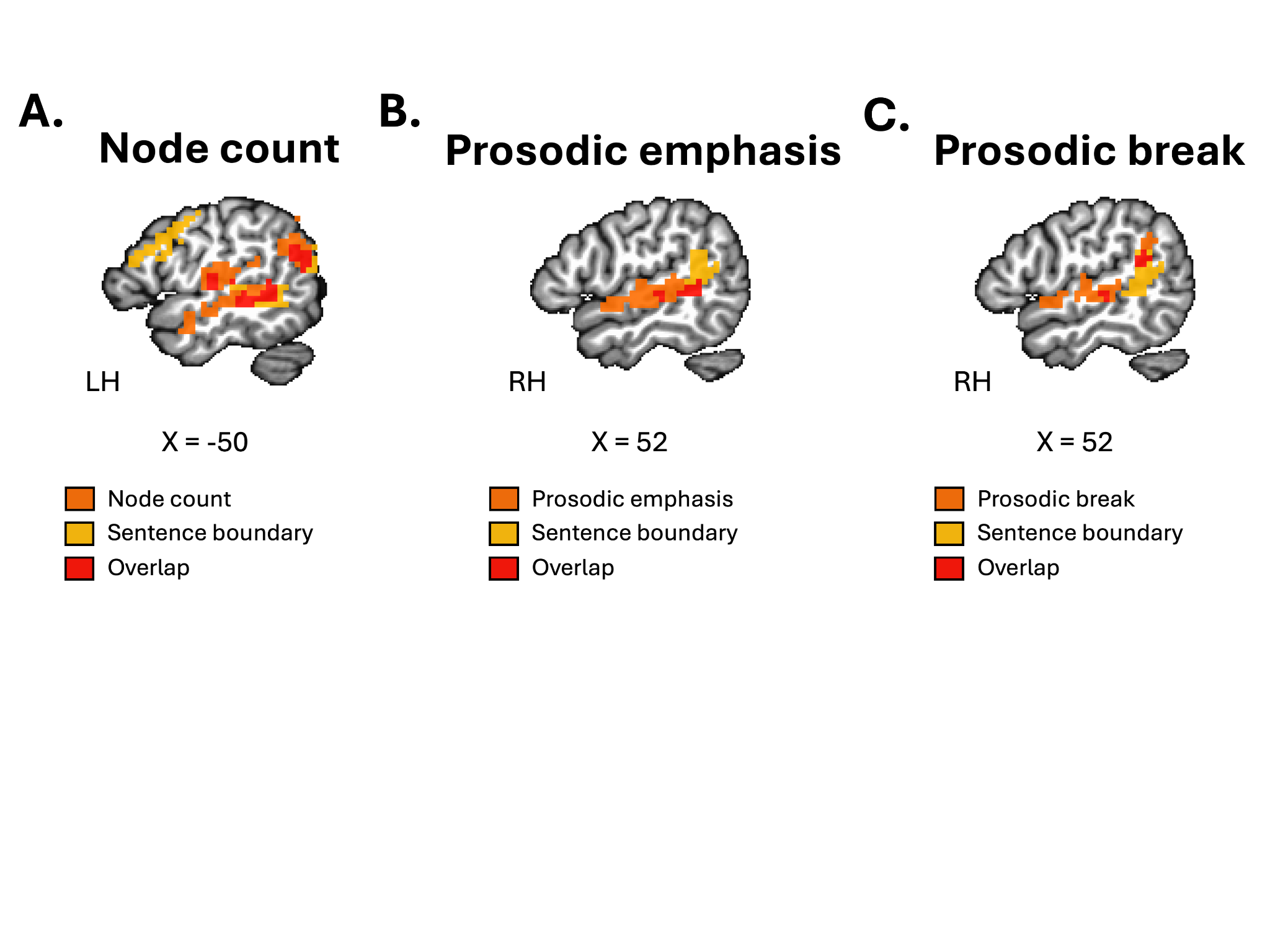

**Figure S1. Spatial overlap between sentence-boundary responses and syntactic and prosodic predictors.** **(A)** Overlap between sentence-boundary and bottom-up node count responses. **(B)** Overlap between sentence-boundary and prosodic emphasis responses. **(C)** Overlap between sentence-boundary and prosodic break responses. Orange indicates significant responses associated with the respective predictor, yellow indicates significant sentence-boundary responses, and red indicates spatial overlap between the two thresholded maps. Cluster masks were extracted after thresholding statistical maps at voxelwise *p* < .01 and cluster-corrected at NN1 with a minimum cluster extent of 39 voxels. Slice coordinates are shown in Talairach TT_N27 space.

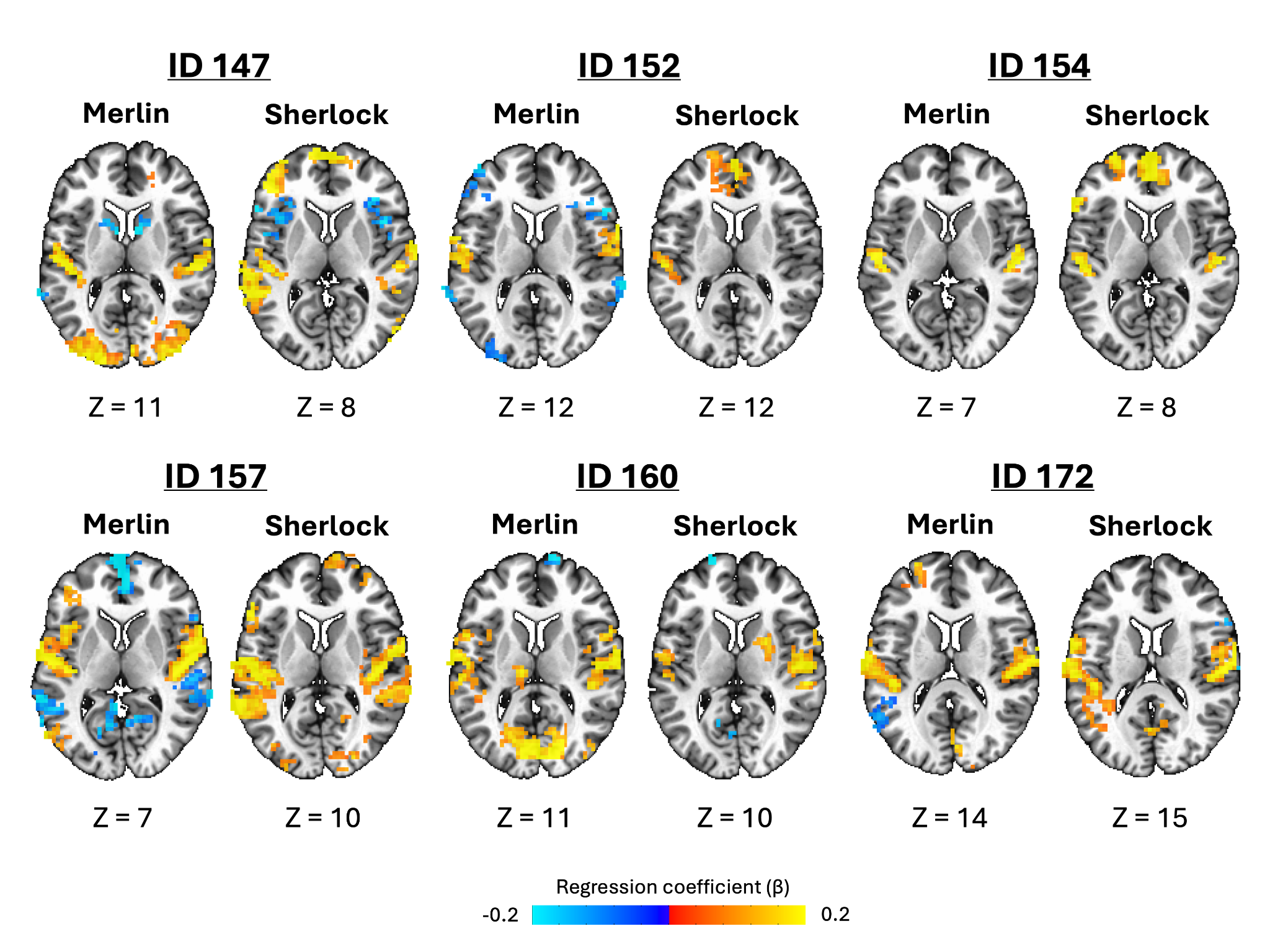

**Figure S2. Individual-participant responses associated with the acoustic envelope across narratives.** Statistical maps are shown for six representative participants for the *Merlin* and *Sherlock* narratives. Warm colors indicate positive effects and cool colors indicate negative effects. Maps are shown as individual-participant regression coefficients (β), thresholded at voxelwise *p* < .01 and cluster-corrected at NN1 with a minimum cluster extent of 39 voxels. Only effects with |β| ≥ 0.2 are displayed. Slice coordinates are shown in Talairach TT_N27 space.
